# *Cryptococcus neoformans* recovered from olive trees (*Olea europaea*) in Turkey reveal allopatry with African and South American lineages

**DOI:** 10.1101/729145

**Authors:** Çağrı Ergin, Mustafa Şengül, Levent Aksoy, Aylin Döğen, Sheng Sun, Anna F. Averette, Christina A. Cuomo, Seyedmojtaba Seyedmousavi, Joseph Heitman, Macit Ilkit

## Abstract

*Cryptococcus* species are life-threatening human fungal pathogens that cause cryptococcal meningoencephalitis in both immunocompromised and healthy hosts. The natural environmental niches of *Cryptococcus* include pigeon (*Columba livia*) guano, soil, and a variety of tree species such as *Eucalyptus camaldulensis, Ceratonia siliqua, Platanus orientalis*, and *Pinus* spp. Genetic and genomic studies of extensive sample collections have provided insights into the population distribution and composition of different *Cryptococcus* species in geographic regions around the world. However, few such studies examined *Cryptococcus* in Turkey. We sampled 388 *Olea europaea* (olive) and 132 *E. camaldulensis* trees from 7 locations in coastal and inland areas of the Aegean region of Anatolian Turkey in September 2016 to investigate the distribution and genetic diversity present in the natural *Cryptococcus* population. We isolated 84 *Cryptococcus neoformans* strains (83 *MAT*α and 1 *MAT***a**) and 3 *Cryptococcus deneoformans* strains (all *MATa*) from 87 (22.4% of surveyed) *O. europaea* trees; a total of 32 *C. neoformans* strains were isolated from 32 (24.2%) of the *E. camaldulensis* trees, all of which were *MAT*α. A statistically significant difference was observed in the frequency of *C. neoformans* isolation between coastal and inland areas (*P* < 0.05). Thus, *O. europaea* trees could represent a novel niche for *C. neoformans*. Interestingly, the *MAT***a** *C. neoformans* isolate was fertile in laboratory crosses with VNI and VNB *MAT*α tester strains and produced robust hyphae, basidia, and basidiospores, thus suggesting potential sexual reproduction in the natural population. Sequencing analyses of the *URA5* gene identified at least 5 different genotypes among the isolates. Population genetics and genomic analyses revealed that most of the isolates in Turkey belong to the VNBII lineage of *C. neoformans*, which is predominantly found in southern Africa; these isolates are part of a distinct minor clade within VNBII that includes several isolates from Zambia and Brazil. Our study provides insights into the geographic distribution of different *C. neoformans* lineages in the Mediterranean region and highlights the need for wider geographic sampling to gain a better understanding of the natural habitats, migration, epidemiology, and evolution of this important human fungal pathogen.

## INTRODUCTION

Cryptococcosis is a potentially lethal disease, especially in immunocompromised hosts, around the world. It is caused by environmental encapsulated yeasts belonging to the *Cryptococcus* genus, including the *C. neoformans* and *C. gattii* species complexes (Hagen et al., 2015; Kwon-Chung et al., 2017). *C. neoformans* has been mainly recovered from pigeon (*Columba livia*) droppings, urban environments, and soil (Lin and Heitman, 2006; May et al., 2016). In addition, it has been isolated from various tree species (Ellis and Pfeiffer, 1990; Randhawa et al., 2008, 2011; Cogliati et al., 2016a, b). Following the first report of *C. neoformans* isolation from trees in Australia (Ellis and Pfeiffer, 1990), many studies have confirmed the environmental association of *Cryptococcus* with plants in different climatic zones (Granados and Castañeda, 2005, 2006; Randhawa et al., 2008, 2011; Bedi et al., 2012; Chowdhary et al., 2012). Several studies have characterized the properties of these yeasts that contribute to the colonization of new environmental niches (Granados and Castañeda, 2006; Randhawa et al., 2008; Ergin and Kaleli, 2010; Ergin et al., 2014; Sengul et al., 2019). With the exception of iatrogenic (Baddley et al., 2011) and zoonotic (Nosanchuk et al., 2000; Lagrou et al., 2005; Singh et al. 2018) cases, *Cryptococcus* infection is caused by the inhalation of airborne basidiospores or desiccated yeast cells from the environment (Hull et al., 2005; Lin and Heitman, 2006; Velagapudi et al., 2009; Springer et al., 2013; May et al., 2016), emphasizing the importance of identifying the natural reservoirs of *C. neoformans* and the molecular links between environmental and clinical isolates and their association with disease (Litvintseva et al., 2005; Noguera et al., 2015; Chen et al., 2015; Kangogo et al., 2015; Spina-Tensini et al., 2017). In a recent study, MLST analysis revealed that some *C. neoformans* genotypes (especially ST63) in Mediterranean countries may be genetically linked (Cogliati et al., 2019). In the environment, the most prevalent mating type is *MATα* (Kwon-Chung and Bennett, 1978).

Most areas colonized by *C. neoformans* are characterized by the presence of several trees, including 4 dominant species: *Eucalyptus camaldulensis* (Mahmoud, 1999; Bernardo et al., 2001; Campisi et al., 2003; Ergin et al., 2004; Gokcen and Ergin, 2014; Mseddi et al., 2011; Romeo et al., 2011, 2012; Colom et al., 2012; Cogliati et al., 2016a, b; Elhariri et al., 2016; Ellabib et al., 2016), *Ceratonia siliqua* (Colom et al., 2012; Romeo et al., 2012; Cogliati et al., 2016a), *Olea europaea* (Cogliati et al., 2016a; Ellabib et al., 2016), and *Pinus* spp. (Cogliati et al., 2016a, b). Further, studies have described numerous woody plants colonized by *C. neoformans* in the Mediterranean region (Mahmoud, 1999; Bernardo et al., 2001; Campisi et al., 2003; Ergin et al., 2004; Mseddi et al., 2011; Romeo et al., 2011, 2012; Colom et al., 2012; Gokcen and Ergin, 2014; Cogliati et al., 2016a, b; Elhariri et al., 2016; Ellabib et al., 2016). *C. neoformans* tree colonization has been observed in northern Mediterranean countries such as Spain (Colom et al., 2012; Cogliati et al., 2016a), Portugal (Bernardo et al., 2001; Ferreira et al., 2014; Cogliati et al., 2016a), France (Cogliati et al., 2016a, b), Italy (Campisi et al., 2003; Romeo et al., 2011, 2012; Cogliati et al., 2016a, b), Greece (Cogliati et al., 2016a, b), and Turkey (Ergin et al., 2004; Ergin, 2010; Ergin and Kaleli, 2010; Gokcen and Ergin, 2014; Cogliati et al., 2016a, b; Sengul et al., 2019), as well as in the northern parts of Cyprus (Cogliati et al., 2016a), Libya (Cogliati et al., 2016a; Ellabib et al., 2016), Tunisia (Mseddi et al., 2011), and Egypt (Mahmoud, 1999; Elhariri, 2016). The olive tree is one of the oldest known cultivated trees in the world and is grown in the entire Mediterranean basin mostly for commercial reasons (Uylaşer and Yildiz, 2014). Although the *Olea* genus is distributed throughout Europe, Asia, Oceania, and Africa, only *O. europaea* is a cultivated species, and recent studies have reported colonization of *O. europaea* with *C. neoformans* in Spain (Cogliati et al., 2016a) and Libya (Ellabib et al., 2016).

In this study, we screened *O. europaea* and *E. camaldulensis* trees in southwestern Anatolia for *Cryptococcus* spp. The isolates we recovered were genotypically diverse, including mating types. Additionally, whole genome sequencing and phylogenomic analyses showed that most of the isolates in Turkey belong to the VNB lineage of *C. neoformans* and are closely related to isolates from Zambia and Brazil. Our studies provide insight into the global distribution, epidemiology, and evolution of this important human fungal pathogen.

## MATERIALS and METHODS

### Study areas

Samples were taken in September 2016 from 7 areas along the Aegean coastal line of Anatolia, Turkey to screen *O. europaea* and *E. camaldulensis* trees for *Cryptococcus* spp. (**Figure 1**). Geospatial characteristics of each sampling area, including climate and geographical coordinates, are presented in **Table 1**, and mean monthly average temperatures and precipitation levels obtained from both local stations and climatic resources (https://sites.ualberta.ca/~ahamann/data/climateeu.html) are shown in **Figure 2**. *E. camaldulensis* (tree symbol in **Figure 1**) is known to be continuously colonized by *C. neoformans* (Ergin et al., 2004; Ergin, 2010) and was sampled to evaluate the current yeast colonization status and mating type distribution.

**FIGURE 1.**
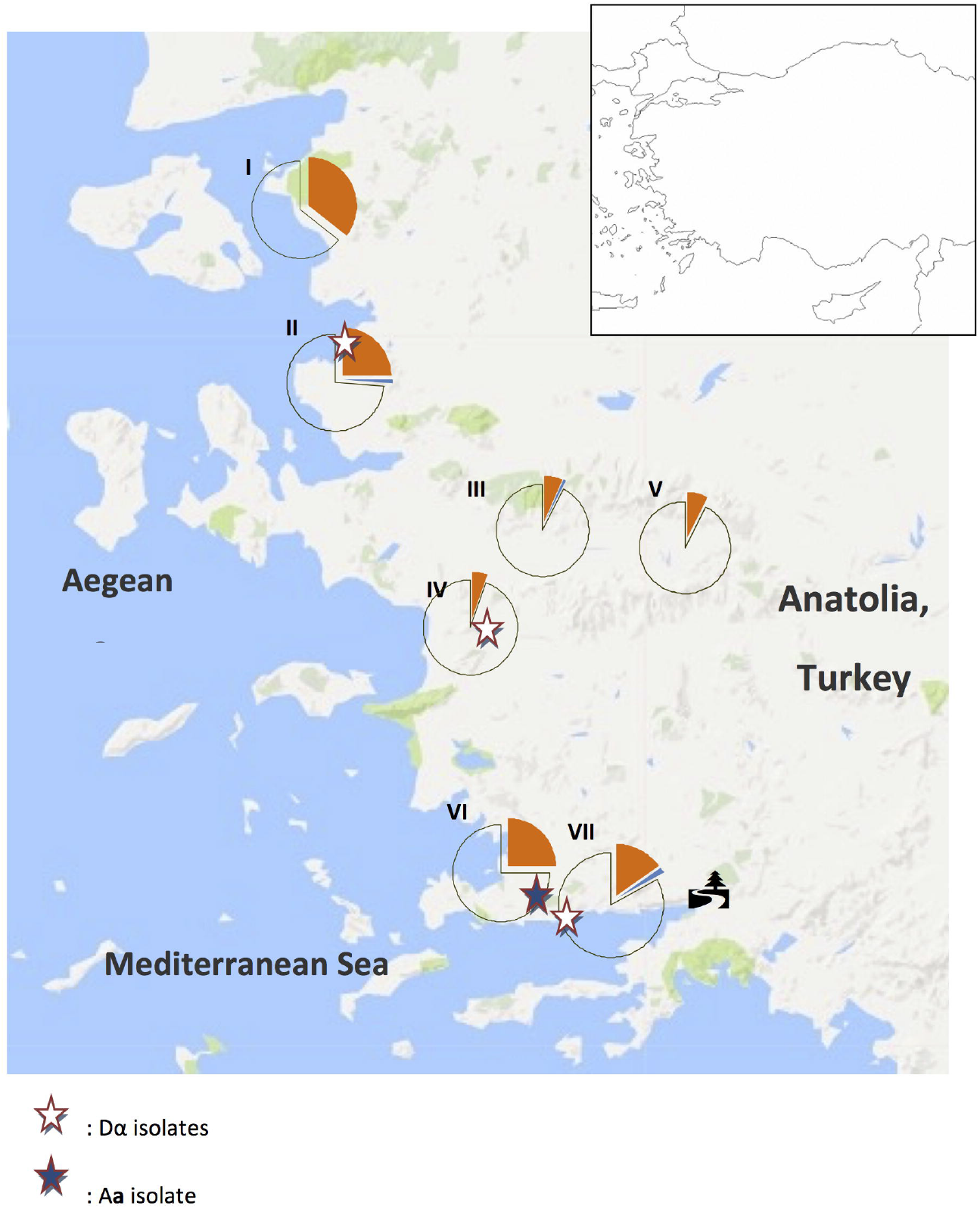
Distribution of trees with *Cryptococcus neoformans* (orange) and *C. deneoformans* (blue) colonization and uncolonized (unshaded) trees in Aegean Anatolia, Turkey. The tree symbol designates the region where *C. neoformans* is recurrently isolated from *Eucalyptus camaldulensis*.

**FIGURE 2.**
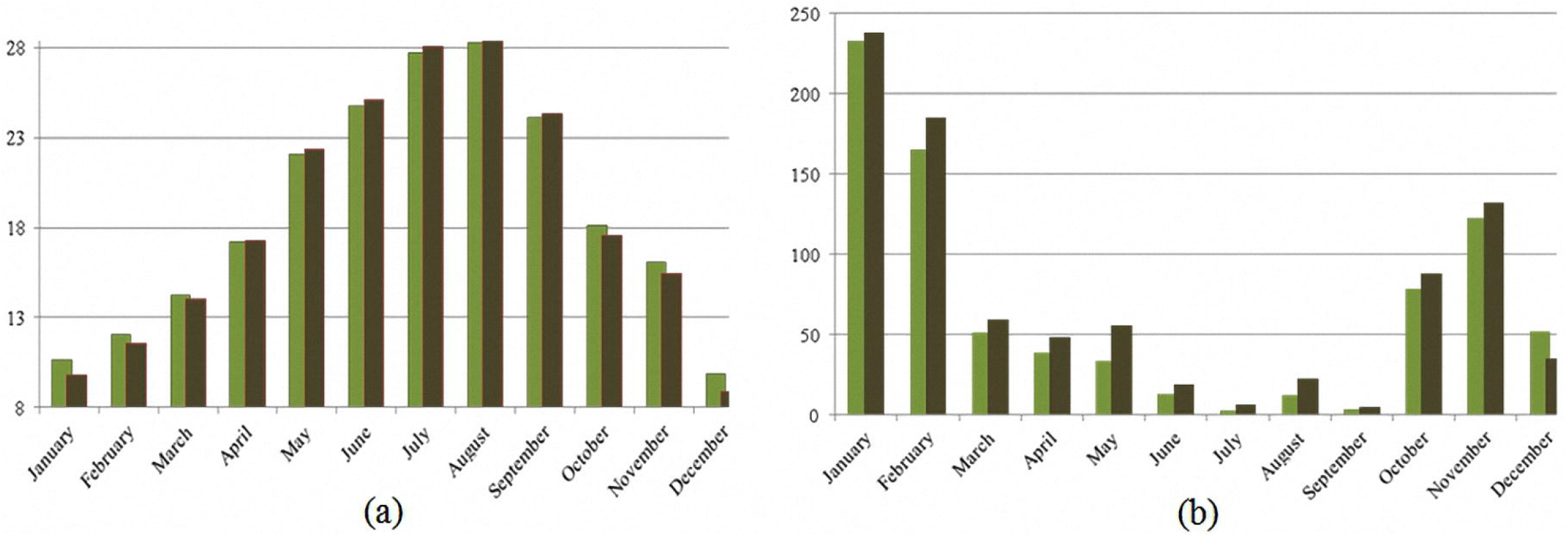
Mean monthly average temperature (a) and precipitation (b) in the sampled coastal regions (light bars) and inland regions (dark bars). Data were obtained in 2016. Mean values are shown.

**TABLE 1.**
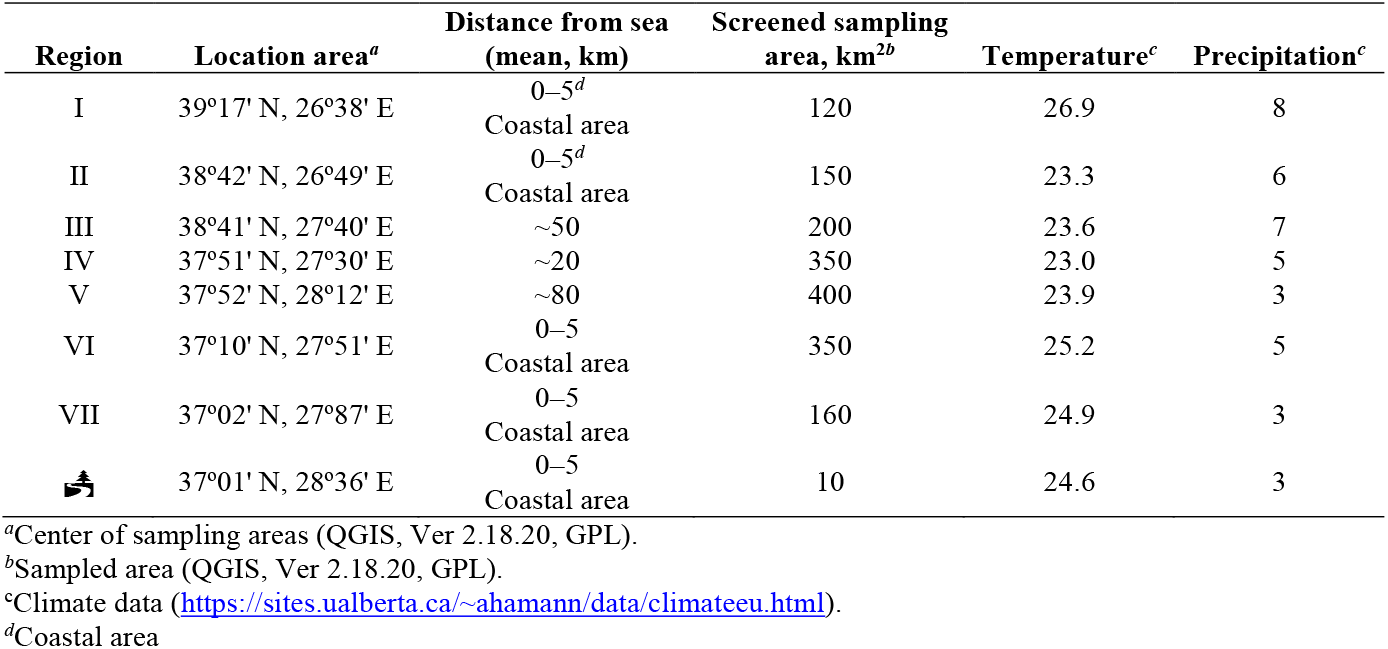
Geographical characteristics of sampling regions.

### Sampling, culture, and conventional identification of *C. neoformans*

In total, 388 typical old *O. europaea* and 132 old *E. camaldulensis* trees from the Mediterranean part of Western Turkey were screened in the study. Tree trunks were randomly sampled by rubbing with a sterile cotton-tipped swab, as described by Randhawa et al. (2005). The swabs were soaked in 3 mL sterile saline containing chloramphenicol (10 mg/L) and transferred to the laboratory at 26°C within 48 h. After vortexing, swabs were removed, and samples were left to sediment for 30 min. Then, undiluted sample supernatants (100 μL) were used to inoculate Staib agar plates containing 0.5% (*w/v*) biphenyl. The plates were incubated at 26°C for 10 d, and moist, characteristically brown-pigmented colonies were analyzed for parameters conventionally used to identify *C. neoformans*, including urea hydrolysis, nitrate reduction, phenoloxidase production, growth at 37°C, and negative reaction on L-canavanine-glycine-bromothymol blue medium (**Table S1**).

### Genomic DNA extraction, polymerase chain reaction (PCR) amplification, and DNA sequencing

All isolates were collected directly from yeast-peptone-dextrose (YPD; Difco, Detroit, MI) agar plates after the second passage. Genomic DNA was extracted using the MasterPure yeast DNA purification kit (Epicentre Biotechnologies, Madison, WI) according to the manufacturer’s instructions.

The species identity and mating type of *C. neoformans* isolates were analyzed by PCR using primers specific for internal transcribed spacer (ITS) and *STE20* genes, respectively (**Table S2**). PCR assays were conducted in a PTC-200 automated thermal cycler in a total reaction volume of 25 μL containing 300 ng of template DNA, 10 pM of each primer, 2 mM of each dNTP, 2.5 μL of 10× Ex Taq buffer, 0.25 μL of ExTaq polymerase (Takara, Shiga, Japan), and an appropriate volume of distilled water. The following cycling conditions were used for PCR with ITS1 and ITS4 primers: initial denaturation at 94°C for 5 min, followed by 36 cycles of denaturation at 94°C for 1 min, annealing at 57°C for 1 min, and extension at 72°C for 1 min, with a final extension at 72°C for 10 min. For PCR with A**a**, Aα, and Dα mating-type primers, cycling conditions were as follows: 95°C for 6 min, followed by 36 cycles at 95°C for 45 s, 60°C for 45 s, and 72°C for 90 s, and a final extension step at 72°C for 6 min. For amplification using D**a** mating-type primers, the cycling protocol was as follows: 95°C for 6 min, followed by 30 cycles at 95°C for 45 s, 50°C for 45 s, and 72°C for 90 s, and a final extension step at 72°C for 6 min. Sterile water was used instead of DNA in negative control samples. Congenic *C. neoformans* strains H99 (VNI-Aα) and KN99**a** (A**a**), and *C. deneoformans* strains JEC20 (VNIV-D**a**) and JEC21 (VNIV-Dα) were used as positive controls in each assay. PCR products were analyzed on 1% agarose gels.

To sequence the ITS region, amplified products were purified using the QIAquick PCR Purification Kit (Qiagen, Germantown, MD) as recommended by the manufacturer. Both DNA strands were sequenced using the BigDye Terminator v. 3.1 cycle sequencing ready reaction mix (Applied Biosystems, Foster City, CA) in an ABI 3130 automated sequencer (Applied Biosystems). Sequences were assembled using the Sequencher 4.8. software (Gene Code Corporation, Ann Arbor, MI).

### Mating assay

Each of the study strains were tested for the ability to mate and compared to the reference *C. neoformans* strains H99 (VNI-Aα) and KN99**a** (A**a**) using mating assays. Mating abilities for the MAT**a** and MATα strains derived from isolate AD215, AD215-D1 and AD215-D2, their mating abilities were tested in crosses between themselves, as well as with the tester strains H99 and Bt3 (VNBI-A**a**). For the mating assay, strains were pregrown on YPD solid medium (Difco) at 30°C for 2 days. Next, a 5-μL suspension of each strain was grown on Murashige Skoog (MS) medium supplemented with a 5-μL suspension of each reference strain with known mating behavior. Cultures were incubated at 25°C in the dark for 2 weeks (Idnurm and Heitman, 2005; Li et al., 2012). Hyphae and basidiospore formation were assessed by light microscopy every other day (**Figure 3**).

**FIGURE 3.**
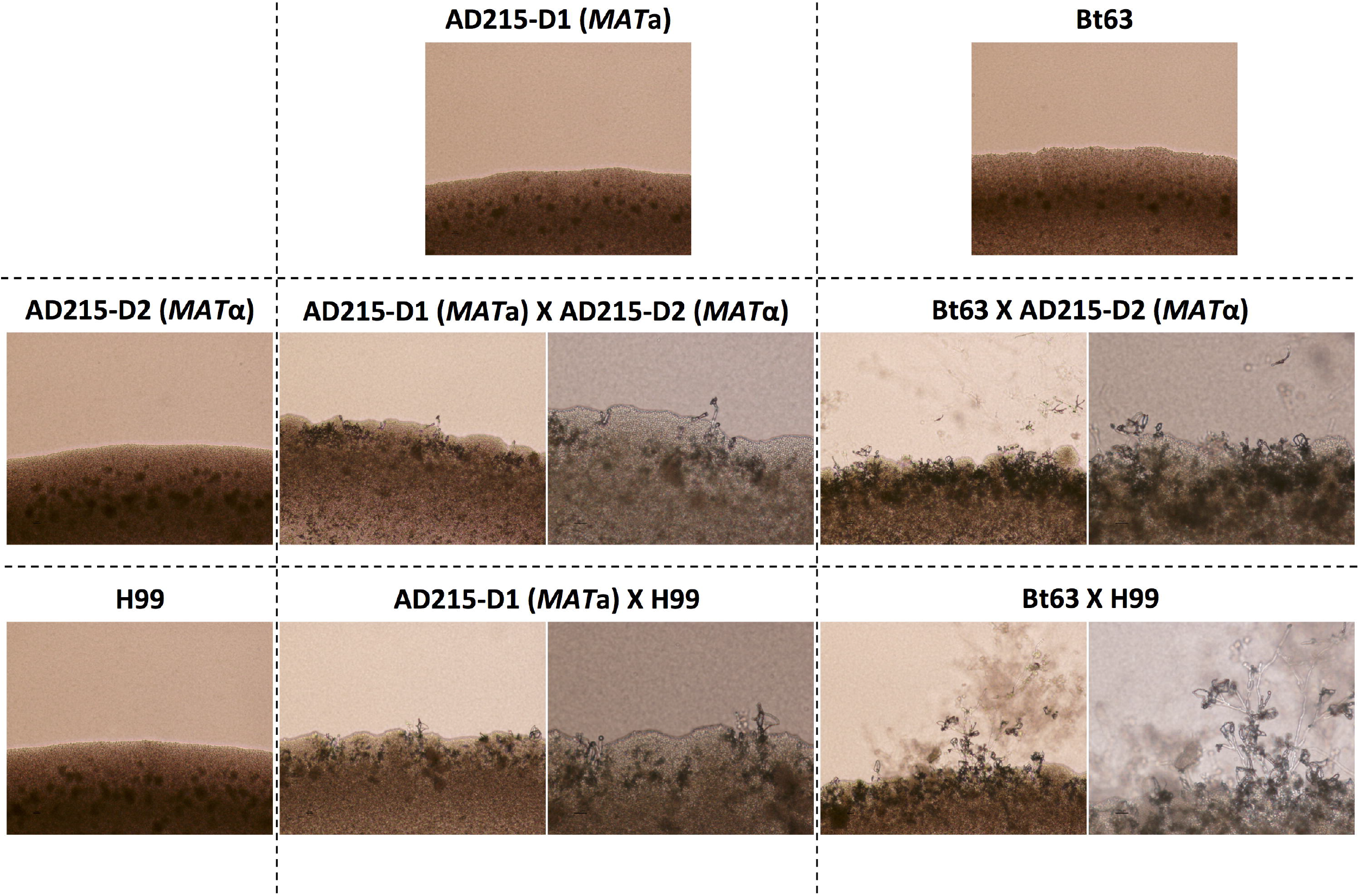
Successful mating between natural *MAT*a and *MAT*α isolates of *Cryptococcus neoformans*. Top: Images of solo cultures of a *MAT***a** colony of strain AD215-D1 and *MAT***a** tester strain Bt63; left: images of solo cultures of a *MAT*α colony from strain AD215-D2 and *MAT*α tester strain H99. All images of solo cultures were taken with 10× magnification. Images of pairwise mating between *MAT***a** and *MAT*α strains are shown within the 2×2 grid: left, 10× magnification; right, 20× magnification. All crosses were carried out on MS medium.

### Genome sequencing and phylogenomic analyses

Whole genome sequences were generated for 9 isolates (AD116, AD119, AD129, AD130, AD131, AD132, AD215-D1, and AD215-D2). Whole genome sequencing libraries were constructed using the Illumina Nextera XT protocol and sequenced on a HiSeqX, generating 150 base-paired end reads (accessible in the NCBI SRA under BioProject PRJNA533587). For SNP calling, reads were downsampled to ~130X sequence depth using Samtools view. Data from a large population survey of 387 isolates (Desjardins et al., 2017), from a Zambian collection (Vanhove et al., 2017) and of VNB isolates from Brazil (Rhodes et al., 2017) were also included. For this set of 446 isolates, reads for each isolate were aligned to the *C. neoformans grubii* H99 assembly (GenBank accession GCA_000149245.2) using BWA-MEM version 0.7.12 (Li, 2013). Variants were then identified using GATK version 3.4 (McKenna et al., 2010). Briefly, indels were locally realigned, haplotypeCaller was invoked in GVCF mode with ploidy = 1, and genotypeGVCFs was used to predict variants in each strain. All VCFs were then combined and sites were filtered using variant filtration with QD < 2.0, FS > 60.0, and MQ < 40.0. Individual genotypes were filtered if the minimum genotype quality was <50, percent alternate allele was <0.8, or depth was <10.

For phylogenetic analysis, the 1,269,132 sites with an unambiguous SNP in at least one strain and ambiguity in ≤10% of strains were concatenated, and insertions or deletions at these sites were treated as ambiguous to maintain the alignment. Phylogenetic trees were estimated using FastTreeDP v 2.1.8 with parameters -gtr and -nt.

### Statistical analysis

Data were analyzed by Chi-square test using the Epi Info^TM^ Stat Calc software (v. 7.2.1.0; Centers for Disease Control and Prevention, Atlanta, GA). A 2-tailed *P* value ≤ 0.05 was considered to indicate statistical significance.

## RESULTS

### Environmental parameters characterizing the sampling regions

In the current study, samples were collected from 7 geographic locations in Anatolia, including 4 coastal regions (I, II, VI, and VII in **Figure 1**) and 3 inland areas (III, IV, and V in **Figure 1**). During the sampling period (September 2016), the mean temperature and humidity were higher in the coastal than in the inland regions (25.3°C vs 23.5°C and 5.5 vs 5.0 mm, respectively).

All 7 sampling regions are located within the natural propagation area of *O. europea* (Uylaşer and Yildiz, 2014) with the typical “macchia” vegetation within the Mediterranean climate (Colom et al., 2012).

### Both mating types are present in natural *C. neoformans* isolates from Turkey

In the Aegean region of Anatolia, *C. neoformans* (n = 84) and *C. deneoformans* (n = 3) were isolated from 22.4% (87/388) of sampled *O. europae* trees. Among them, 95.4% (83/87), 1.1% (1/87), and 3.5% (3/87), were identified as serotypes A*MAT*α, A*MAT***a**, and D *MAT*α, respectively (**Figure 4** and **Table 2**), whereas strains of serotype D *MAT***a** were not identified. This corresponded to a significantly higher frequency of *C. neoformans* strains isolated from the beach/coastal regions (75/221) compared to inland areas located ≥10 km from the sea (12/167) (P<0.001; **Table 3**).

**FIGURE 4.**
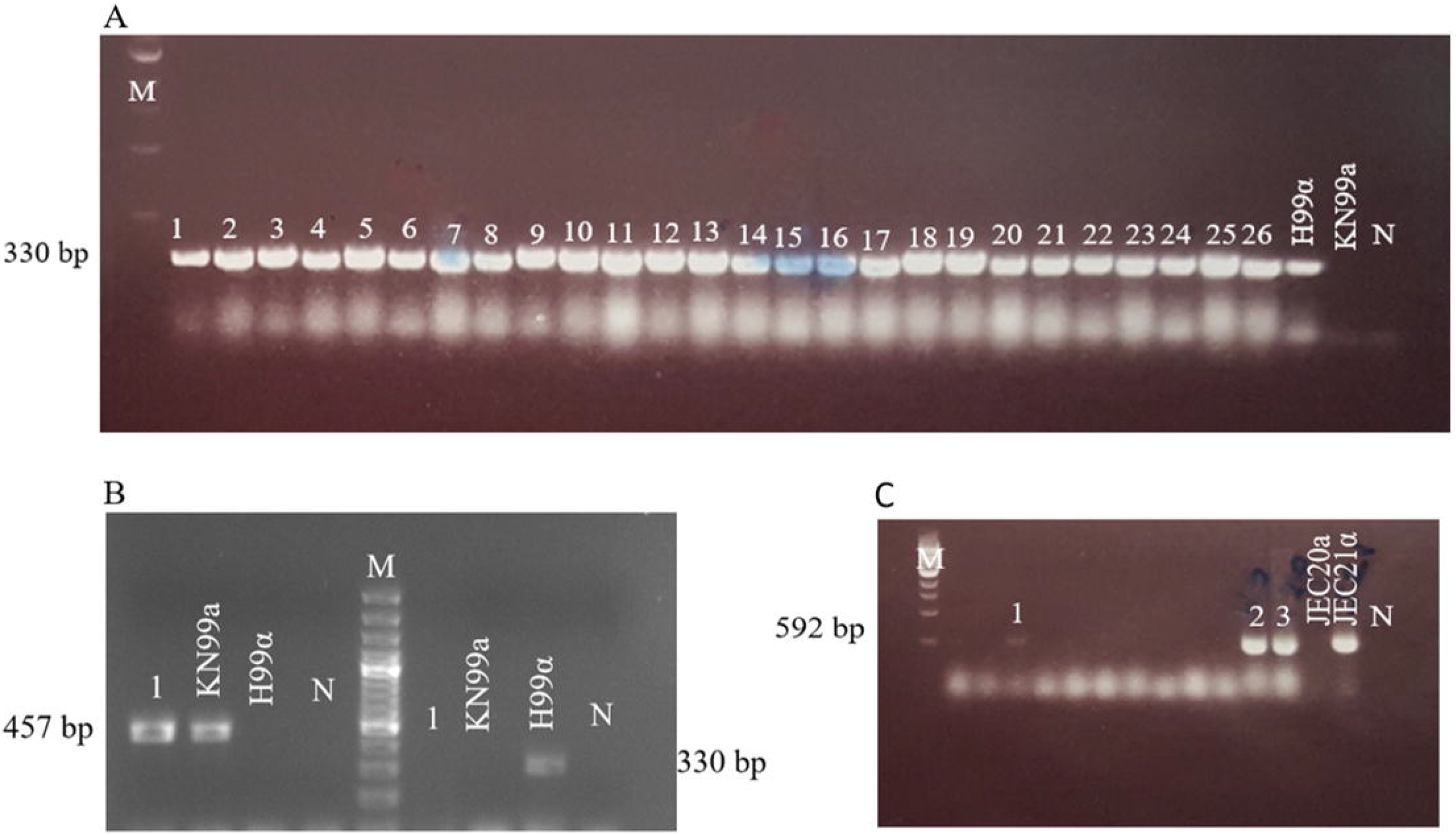
Mating type profiles obtained by PCR using *STE20* gene-specific primers. (A) 1–26: serotype Aα, H99α, KN99**a**; (B) 1: serotype A**a** (strain AD215), KN99**a**, H99α; (C) 1–3: serotype Dα, JEC20**a**, JEC21α. Congenic *C. neoformans* strains H99 (VNI-αA) and KN99**a** (**a**A), and *C. deneoformans* JEC20 (VNIV-**a**D) and JEC21 (VNIV-αD) were used as positive controls. M, molecular weight markers; N, negative control.

**TABLE 2.**
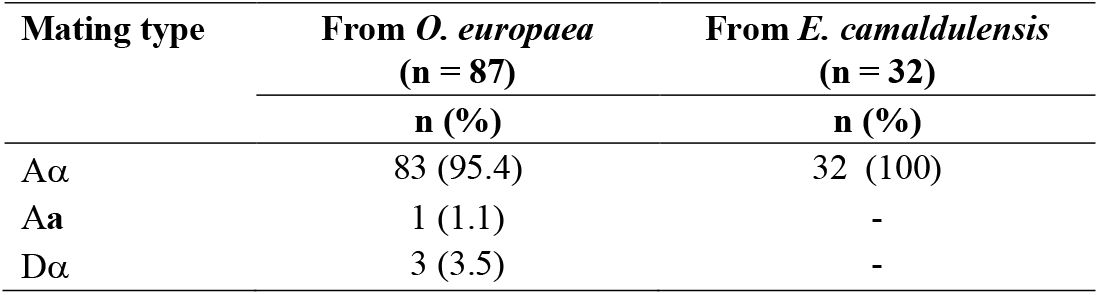
Mating types of *Cryptococcus neoformans* and *C. deneoformans* according to tree species.

**TABLE 3.**
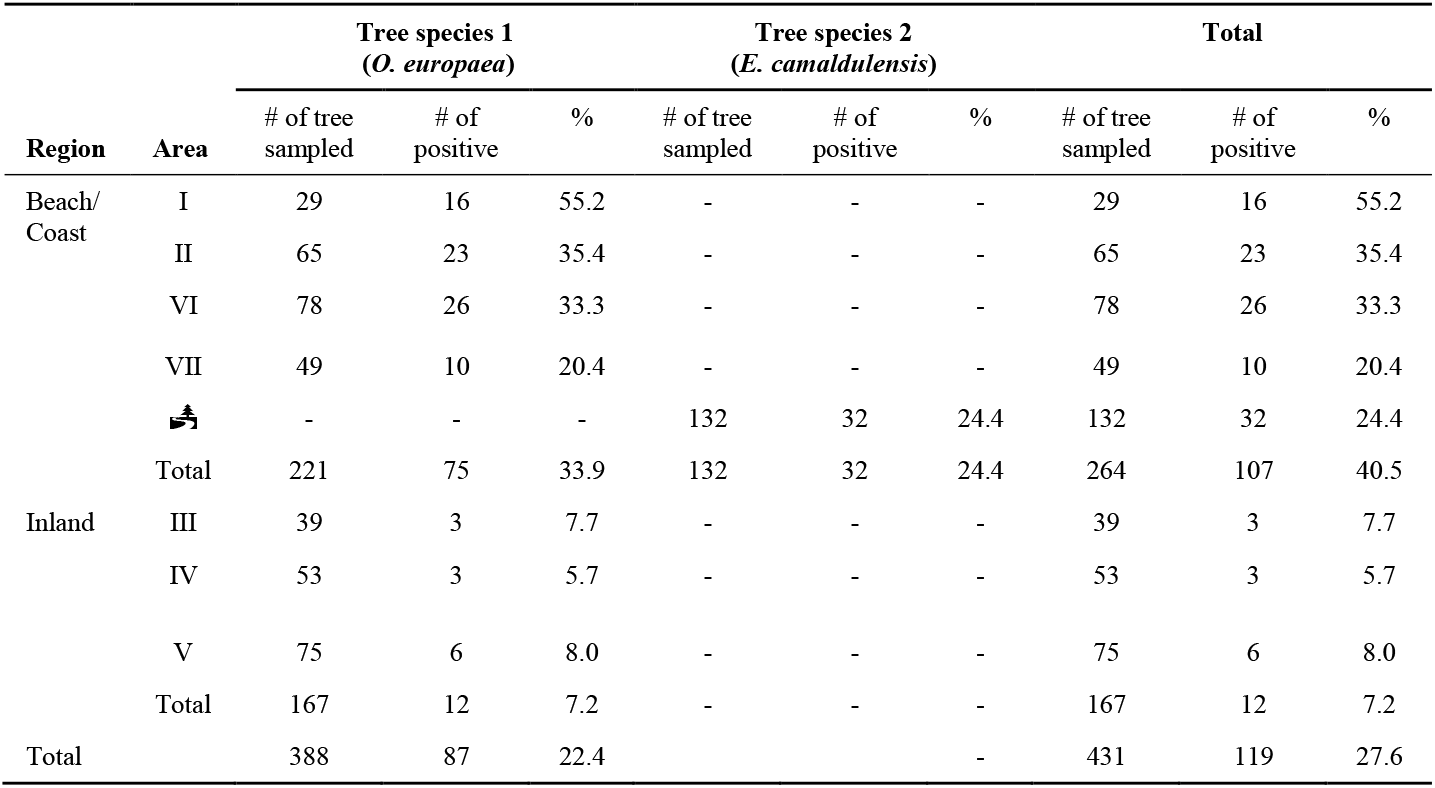
Geographical distribution of *Olea europaea* and *Eucalyptus camaldulensis* colonized with *Cryptococcus neoformans*.

The single *MAT***a** type *C. neoformans* strain (AD215) isolated from area VII showed no sign of mating when crossed with a *MAT***a** tester strain (KN99**a**), but did show signs of robust sexual reproduction when crossed with the *MAT*α tester strain (H99) (**Figure 3**), phenotypically confirming that this isolate is a true *MAT***a** *C. neoformans* isolate.

Thirty-two of 132 (24.2%) isolates have been isolated from *E. camaldulensis* trees, all of which were identified as *MAT*a *C. neoformans*.

### Natural *C. neoformans* isolates in Turkey are genetically diverse (*URA5*)

Because the genotyping of the *MAT* locus showed that the vast majority of the isolates in this study are *MAT*α, we further investigated how much genetic diversity is present within the Turkish *Cryptococcus* isolates. We randomly picked 41 strains and PCR amplified and Sanger sequenced the *URA5* locus, which in previous studies has been suggested to be more genetically diverse among natural *Cryptococcus* strains. Our sequencing analyses identified 5 *URA5* alleles in the 41 isolates (**Figure S1**), including 2 major genotypes represented by 14 and 18 isolates, respectively, 1 genotype represent by 7 isolates, and 2 unique genotypes each represented by 1 isolate. Thus, ample genetic diversity is present among the natural isolates in Turkey.

### Natural *C. neoformans* isolates in Turkey are closely related to those from Brazil and Zambia

To investigate how isolates from Turkey compare to other global *C. neoformans* isolates, we carried out whole genome sequencing for 7 isolates that represent the 5 *URA5* genotypes (**Figure S1**), as well as the *MAT***a** (AD215-D1) and *MAT*α (AD215-D2) strains derived from isolate AD-215, and then compared their genome sequences with those of the global *C. neoformans* sequences that have been recently published (Desjardins et al., 2017, Vanhove et al., 2017) based on variants identified compared to the H99 reference genome.

*URA5* sequences extracted from the variant calls of these isolates are in overall agreement with the *URA5* genotyping results described above. Phylogenetic analysis suggested that 3 of the 7 isolates are VNI (AD119, AD129, and AD130), while the remaining 4 isolates (AD116, AD131, AD132, and AD161) and the 2 strains derived from AD215 (AD215-D1 and AD215-D2) are VNB. The 3 VNI isolates (AD119, AD129, and AD130) were placed within the VNIb subclade of global isolates (**Figure S2**), and all VNB isolates were placed within the VNBII clade, which primarily includes South African isolates (**Figure 5**). Of the VNBII isolates from Turkey, AD116 belongs to a clade that contains mostly isolates from Botswana, South Africa, and Zambia, while the others are more divergent from most VNBII isolates and are closely related to 2 isolates from Brazil and 3 isolates from Zambia (**Figure 5**).

**FIGURE 5.**
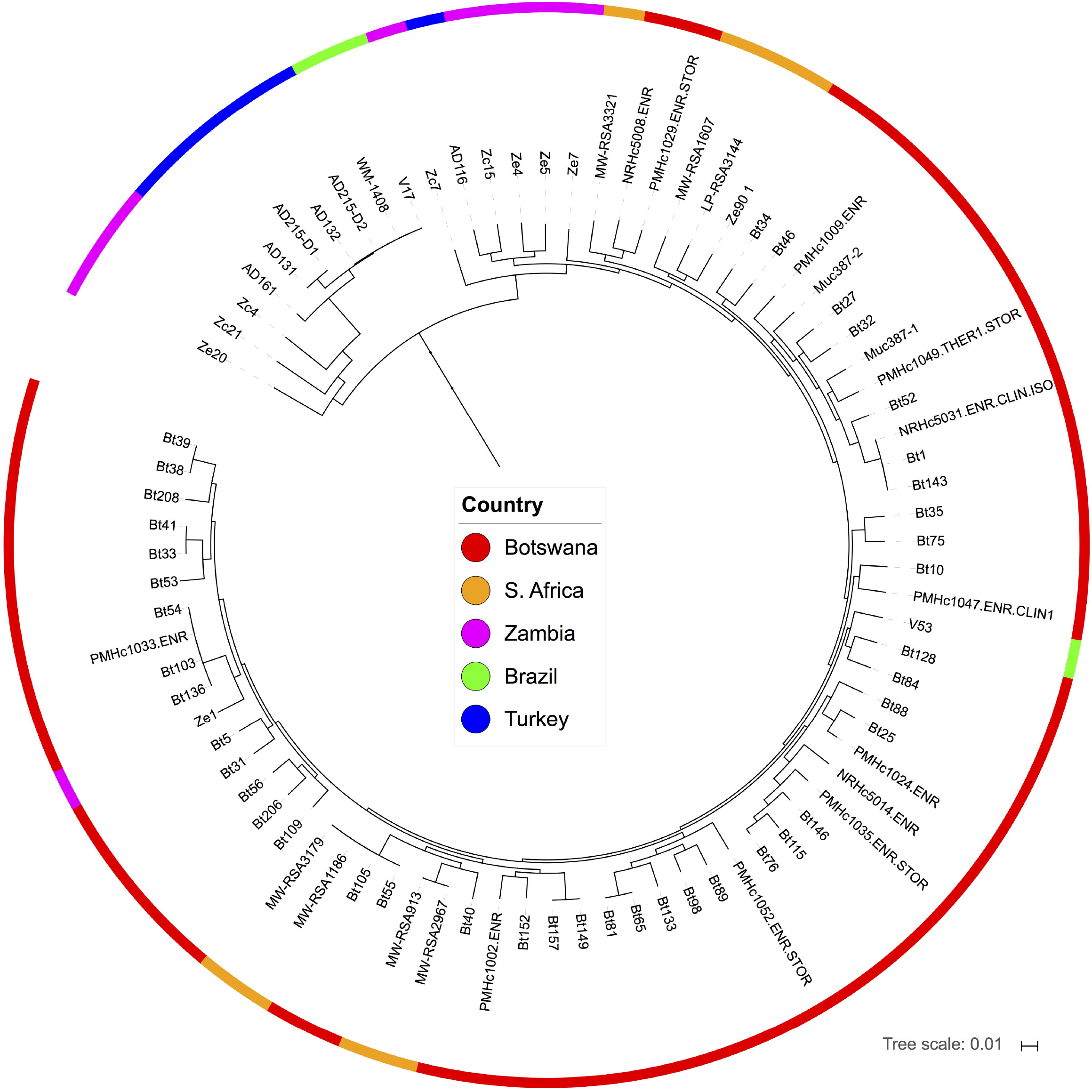
Phylogenic analyses places isolates from Turkey in the VNBII lineage of *C. neoformans*. Isolates from Turkey are part of a divergent subclade that included isolates from Brazil and Zambia. The phylogeny was estimated from 1,269,132 segregating sites using FastTree (Price et al., 2009), and the tree was rooted with VNII as the outgroup.

Interestingly, 2 isolates from Turkey (AD132 and AD215-D2) are separated by only an average of 590 SNPs from 2 isolates from Brazil (an environmental isolate, WM-1408, and a clinical isolate, V17; **Figure 5**). This number of SNPs does not suggest a recent transmission event between these locations, but rather a closer than expected relationship across continents in a lineage formerly thought to be confined to southern Africa.

## DISCUSSION

The present study revealed that olive trees are a major reservoir of environmental *C. neoformans* strains in the Aegean part of the Mediterranean region, where cultivation of olive trees is a tradition (Uylaşer and Yildiz, 2014). We sampled only old tree trunks with hollows that constitute an appropriate yeast habitat, providing stable humidity and temperature and protection from solar radiation (Lin and Heitman, 2006; Velagapudi et al., 2009; May et al., 2016). Compared to inland regions, the colonization of olive trees by *C. neoformans* is more prominent in the coastal areas, which are less arid, are at a lower altitude, and have milder winters. Throughout the study period (September 2016), the mean temperatures and precipitation in the coastal areas were higher than those in the inland areas, and accordingly, the largest number of *C. neoformans* isolates was obtained from area I, which had the highest temperature and precipitation (**Tables 1** and **3**; *P* < 0.01).

In the current study, *C. deneoformans* was obtained from areas II, IV, and VII. In 2008, *C. deneoformans* was isolated mainly from pigeon droppings in the Aegean region (Karaca Derici and Tümbay, 2008), which corresponds to area II in the current study. However, clinical cases of *C. deneoformans* were documented in the Black Sea coastal area (Kaya et al., 2012; Birinci et al., 2016), which is distant from the Aegean coast and where the climate is more humid and the temperature lower. This finding is consistent with reports that *C. deneoformans* is more sensitive to heat than *C. neoformans* (Martinez et al., 2001; Lin and Heitman, 2006; Bedi et al., 2012). The discrepancy between the environmental presence of *C. neoformans* and related clinical cases should be addressed by further studies screening different areas in Turkey. The northern and southern areas of the Turkish Mediterranean coast have different climatic conditions and, consequently, distinct tree populations, which may potentially influence the rate of cryptococcal colonization.

The first isolation of *C. neoformans* from *E. camaldulensis* in Anatolia was reported in 2004 (Ergin et al., 2004). A previous study investigated the presence of *C. neoformans* in the flowers of *E. camaldulensis* trees from the Akyaka/Gökova (tree symbol, **Figure 1**) districts in southeastern Turkey and identified only 1 *C. neoformans* isolate (0.09%) (Ergin et al., 2004). In 2010, repeated screening of 17 *E. camaldulensis* trees with large trunks was performed in the same region, and a colonization rate of 64% was reported (Ergin, 2010). In the current study, we sampled a significantly higher number of *E. camaldulensis* trees (n = 132) and observed a lower colonization rate of 24.2%, which is probably more representative due to the larger area of analysis.

Trees play an essential role as a reservoir and breeding ground for the propagation of *C. neoformans*. A recent study showed that *Cryptococcus* has the ability to colonize some plant such as *E. camaldulensis*, *Terminalia catappa*, *Arabidopsis thaliana*, *Colophospermum mopane*, *Tsuga heterophylla*, and *Pseudotsuga menziesii*, as well as their debris, which constitutes the ecological niche and reservoirs of infectious propagules of *Cryptococcus* in the environment (Springer et al., 2017). Climate conditions, including humidity, temperature, evaporation, and solar radiation, play significant roles in the environmental distribution of *C. neoformans* (Lin and Heitman, 2006; Velagapudi et al., 2009; Springer et al., 2013; May et al., 2016). Our findings indicate that *C. neoformans* colonization of olive trees reflects the Mediterranean ecological model influenced by climate changes and urbanization (Garcia-Mozo et al., 2016). Several studies reported the presence of environmental *C. neoformans* in Mediterranean countries, including Spain (Colom et al., 2012; Cogliati et al., 2016a) and Libya (Ellabib et al., 2016), based on the association between the climate and yeast distribution. Warmer temperatures can affect *Cryptococcus* spp. spread, especially that of *C. gattii*, a sibling species of *C. neoformans* (Granados and Castañeda, 2006; Randhawa et al., 2011; Bedi et al., 2012; Chowdhary et al., 2012, Uejio et al., 2015; Cogliati et al., 2016b). Although it is seen that the olive trees practiced in the Mediterranean “macchia” are more interrelated with the *C. gattii, C. neoformans* colonization is not far from the ecosystem (Cogliati et al., 2017). The duration of the dry season, natural degradation of woods, and drier habitats account for lower bacterial presence and less competition for nutrients, thus constituting favorable conditions for *C. neoformans* colonization, especially of old tree trunks (Ruiz et al., 1981; Granados and Castañeda, 2006; Cogliati et al., 2017).

Worldwide screening for the presence of *C. neoformans* in the environment has been performed since the mid −1990s, and *E. camaldulensis* was established as the major source of tree-associated cryptococcosis in the early 2000s (Ellis and Pfeiffer, 1990; Campisi et al., 2003; Ergin et al., 2004; Lin and Heitman, 2006; Randhawa et al., 2008; Nougera et al., 2015). Several studies reported that the majority of *C. neoformans* environmental isolates contain *MAT*α mating-type alleles (Litvintseva et al., 2005; Nielsen et al., 2005; Lin and Heitman, 2006; 2014; Chen et al., 2015; Kangogo et al., 2015), which is consistent with our findings that *Cryptococcus* isolates from *E. camaldulensis* have the *MAT*α phenotype. However, in *O. europaea*, we isolated *C. neoformans* harboring Dα and even A**a** alleles. To the best of our knowledge, it is the first *MAT***a**-containing serotype A strain isolated from the environment in Anatolia. A previous study performed by Saracli et al. (2006) in Anatolia identified *C. neoformans* with mating types Aα (65.4%), D**a** (15.4%), and Dα (11.5%), but not A**a**, in pigeon droppings. Although *MAT*α predominates in clinical and environmental populations, 4 *C. neoformans MAT***a** VNI isolates (125.91, IUM96–2828, Bt130, and IUM99–3617) were previously identified from both clinical and environmental sources (Lengeler et al., 2000; Viviani et al., 2001, 2003; Keller et al., 2003; Nielsen et al., 2003; Litvintseva et al., 2007). Whole genome analysis of 3 of these isolates (125.91, IUM96–2828, and Bt130) suggests introgression of *MAT***a** into VNI strains (Desjardins et al., 2017; Rhodes et al., 2017). In the present study, isolation of a novel serotype A *MAT***a** strain from the environment suggests that **a**-α sexual reproduction might occur in the serotype A population. However, whole genome analyses of additional isolates is required to detect recombination signatures in the Turkish *Cryptococcus* population.

*Cryptococcus* isolates in Turkey are genetically diverse. Based on our *URA5* genotyping and whole genome sequencing analyses, most isolates from Turkey belong to the VNI and VNBII groups. Specifically, of the 41 isolates that were genotyped for the *URA5* locus, 21 likely belong to the VNI group that includes isolates AD119, AD129, and AD130, while the other 20 belong to the VNBII group that includes isolates AD116, AD131, AD132, and AD161. Interestingly, of the VNBII isolates, only 1 (AD116) belongs to the larger sublineage that contains most VNBII isolates from Botswana, South Africa, and Zambia, while the other 19 have *URA5* alleles identical to isolates AD131, AD132, and AD161, which, based on whole genome sequence analyses, belong to within the smaller VNBII sublineage that includes strains from Botswana and Brazil. Two strains from Brazil (V17 and WM-1408) may have contributed significant genetic material to the other lineages (VNI, VNII, and VNB) through recombination, with V17 donating the most genetic material to VNI isolates in Africa, India, and Thailand (Rhodes et al., 2017). The isolation of Turkish isolates, including the *MAT***a** and *MAT*α fertile strains derived from isolate AD215, that are almost genetically identical to V17 and WM-1408, suggests that it is possible that the Mediterranean region could be a fertile ground for genetically diverse *Cryptococcus* isolates and could serve as an important center for the global migration and distribution of *Cryptococcus* isolates.

In conclusion, our results indicate that compared to *C. deneoformans, C. neoformans* is more common on olive trees and *E. camaldulensis* in the Aegean region of Anatolia. While the vast majority of the natural isolates in Turkey are mating type α, the presence of a fertile *MAT***a** isolate suggests that sexual reproduction could be ongoing in natural *C. neoformans* populations. Our finding that *C. neoformans* isolates from Turkey belong to VNBII and are more closely related to strains from Zambia and Brazil provides insights into the global distribution of *C. neoformans* and emphasizes the need for more extensive environmental screening to reveal new reservoirs for *C. neoformans*, which would promote our understanding of the natural distribution, epidemiology, and evolution of this important human fungal pathogen.

## DATA AVAILABILITY

The data analyzed in this study are accessible at NCBI SRA under BioProject PRJNA533587.

## AUTHOR CONTRIBUTIONS

*Conceived and designed the experiments:* ÇE, AD, SS, CC, JH, and MI. *Performed the experiments:* ÇE, MŞ, LA, AD, SS, AA, and CC. *Analyzed the data:* ÇE, AD, SS, CC, JH, and MI. *Contributed reagents/materials/analysis tools:* ÇE, AD, SS, CC, JH. *Drafted and revised the manuscript:* ÇE, MS, LA, AD, SS, AA, CC, SS, JH, and MI. All authors have read, revised, and approved the final manuscript.

## ACKNOWLEDGMENTS

The authors are grateful to the members of the Heitman laboratory at the Duke University Department of Molecular Genetics and Microbiology (Durham, NC, USA) for their valuable assistance with laboratory analyses.

## ETHICAL APPROVAL

This article does not contain any experiments involving human participants or animals.

## CONTRIBUTION TO THE FIELD STATEMENT

*Cryptococcus neoformans* is one of major human fungal pathogens. Studies of their natural distribution, existing genetic diversity, and population dynamics contribute to our better understanding of the epidemiology and the evolution of the virulence in these fungal pathogens. In this study, we surveyed a large number of *Olea europea* (olive) and *E. camaldulensis* trees for the presence of *Cryptococcus* species from 7 locations that represent the coastal and inland areas of the Aegean Turkey in Turkey. We found that *Cryptococcus* is prevalent in Turkey, and the vast majority of the isolates are *C. neoformans*. There are ample genetic diversity existing among the Turkey *C. neoformans* isolates, with the presence of both *MATa* and *MAT*α strains, suggesting ongoing sexual reproduction among the natural isolates. Interestingly, several of the *C. neoformans* isolates from Turkey show high genetic similarity across the genome with strains isolated from Zambia and Brazil. Our study provides insights into the natural niches and distribution of different *C. neoformans* lineages in the Mediterranean region and helps us to gain a better understanding of the ecology, epidemiology, and evolution of this important human fungal pathogen.

## SUPPORTING INFORMATION

**TABLE S1. Primers used in this study.**

**TABLE S2. *Cryptococcus* isolates recovered and analyzed in this study.**

**FIGURE S1. Genotyping of the *URA5* locus revealed that the isolates from Turkey are genetically diverse.**

**FIGURE S2. Phylogenetic analysis of isolates from Turkey with a diverse set of other global isolates**. The full phylogeny estimated for Figure 5 is shown, including VNI, VNII, VNBI, and VNBII lineages. The phylogeny was estimated from 1,269,132 segregating sites using FastTree (Price et al., 2009), and the tree was rooted with VNII as the outgroup. Isolates from Turkey were placed with the VNIb sublineage of VNI and in VNBII.

