## Supplementary figures and images for "*Cryptococcus neoformans* recovered from olive trees (*Olea europaea*) in Turkey reveal allopatry with African and South American lineages"

### Supp.Fig.1

#### Slide 1
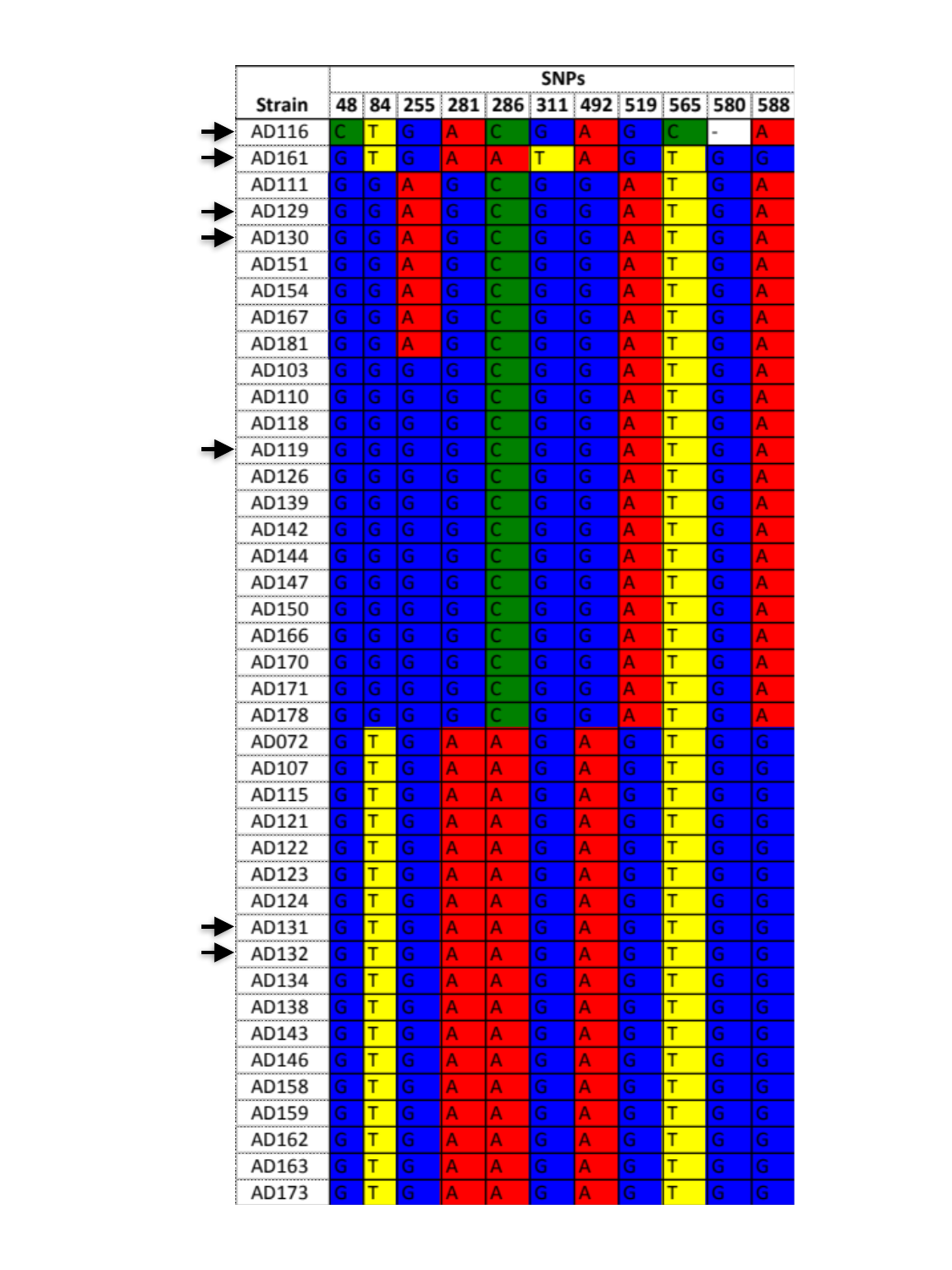

### Supp.Fig.2

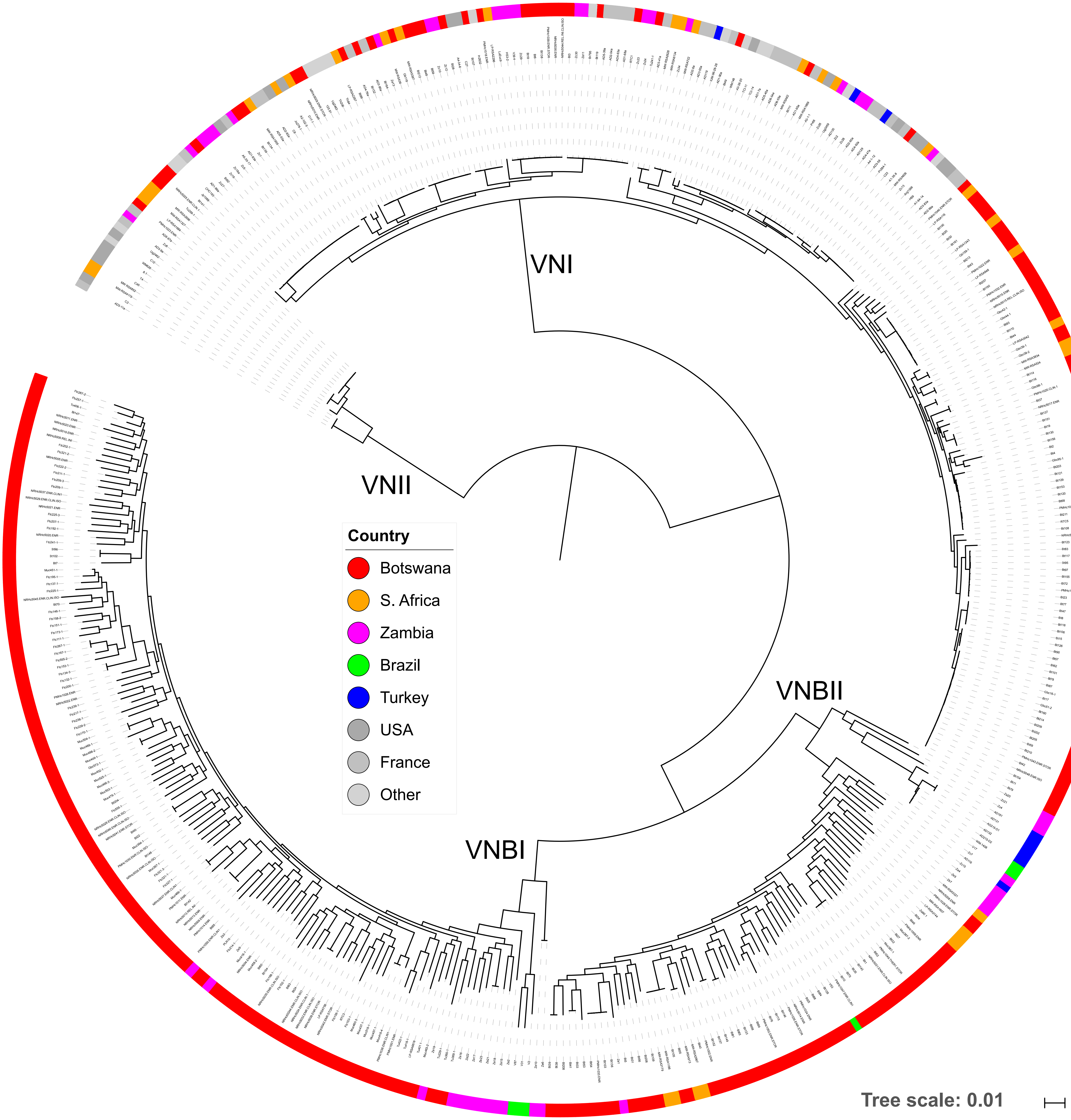
