## Supplementary material for "*Cryptococcus neoformans* recovered from olive trees (*Olea europaea*) in Turkey reveal allopatry with African and South American lineages": Supp.Table2

**S2 Table. Primers used in this study.**

| Target | Primer symbol | Sequence (5′–3′) |
| --- | --- | --- |
| ITS | | |
|  | ITS1  ITS4 | TCCGTAGGTGAACCTGCGG  TCCTCCGCTTATTGATATGC |
| Mating-type alleles | | |
| *C*. *neoformans* | | |
|  | *STE20* A**a** F  *STE20* A**a** R  *STE20* Aα F  *STE20* Aα R | CTAACTCTACTACACCTCACGGCA  CGCACTGCAAAATAGATAAGTCTG  GGCTGCAATCACAGCACCTTAC  CTTCATGACATCACTCCCCTAT |
| *C*. *deneoformans* | | |
|  | *STE20* D**a** F  *STE20* D**a** R  *STE20* Dα F  *STE20* Dα R | CACATCTCAGATGCCATTTTACCA  AGCTCTAAGTCATATGGGTTATAT  CTTAATTCACAGCACCAGCCTA  GGTCATCACAGTCAGTCACCAC |

F, forward; R, reverse.
